# Sex differences in concentrations of HMGB1 and numbers of pigmented monocytes in infants with malaria

**DOI:** 10.1101/2021.03.26.437109

**Authors:** Bernard N. Kanoi, Thomas G. Egwang

## Abstract

Females generally mount more robust innate and adaptive immune responses and demonstrate a higher rate of morbidity, and prevalence of autoimmune diseases by comparison with males. In malaria, females demonstrate higher concentrations of antibodies and rates of severe adverse events and mortality following natural infections and malaria vaccination. Although monocytes/macrophages play a crucial role in disease and protection in malaria, no studies have investigated sex differences in their functions in production of proinflammatory cytokines and chemokines in malaria-infected subjects. Here, we show significant sex differences in serum concentrations of HMGB1, a non-histone chromatin-associated protein, and numbers of pigmented monocytes, which are both markers of severe malaria, in infants <5 years old from a malaria endemic region in Northern Uganda. Female infants with clinical malaria had significantly higher HMGB1 concentrations than male infants, and female infants with asymptomatic malaria had significantly lower numbers of pigmented monocytes than male infants with asymptomatic malaria. There was (1) a significant correlation between HMGB1 concentrations and pigmented monocyte numbers in female but not male infants; and (2) a significant correlation between HMGB1 concentrations and parasite densities in female but not male infants. These findings suggest that female infants with clinical malaria might be at a greater risk of morbidity characterized by higher serum HMGB1 levels.

## Introduction

Malaria causes mortality of close to half a million people annually, mostly in sub-Saharan Africa (1). Reports infer that this mortality is significantly higher in females than males (2). Females generally mount more robust innate and adaptive immune responses and demonstrate a higher rate of morbidity and prevalence of autoimmune diseases by comparison with males (3, 4). Females have demonstrated higher levels of antibodies following natural infections (5) and malaria SPf66 vaccination (6, 7), and higher rates of severe adverse events and mortality following SPf66 and RTS,S/AS01 malaria vaccinations (4, 8). Furthermore, female neonates exposed to malaria *in utero* demonstrate higher numbers of regulatory T-cell (Tregs) at birth (9) and female children clear asymptomatic malaria faster than male children (10).

Although monocytes/macrophages play a crucial role in disease and protection against malaria (11), no studies have investigated sex differences between malaria-infected or exposed individuals in terms of monocyte/macrophage functions such as production of pro and anti-inflammatory cytokines, chemokines and high mobility group box 1 (HMGB1). The characterization of sex differences in mechanisms underlying monocyte/macrophage-mediated protective immunity and morbidity will inform the development of sex-based approaches which reduce morbidity in females and increase antibody responses in males following vaccination. This is key to developing interventions that are equally effective for both sexes.

HMGB1 is a non-histone chromatin-associated protein that is passively released by dead or dying cells as a result of tissue injury following infection with pathogens and therefore serves as a danger signal (12). As a damage associated molecular pattern (DAMP) molecule, extracellular HMGB1 exacerbates inflammation by activating monocytes/macrophages to produce proinflammatory cytokines which characterize severe disease in sepsis, COVID-19, and severe and complicated malaria (13–15). HMGB1 is also actively secreted by activated monocytes/macrophages (16). Both pigmented monocytes and HMGB1 have been characterized as markers of severe malaria in children in malaria endemic regions (17, 18). Here, we present for the first-time data showing significant sex differences in concentrations of HMGB1 and numbers of pigmented leucocytes and their correlations with parasite densities in female and male infants <5 years old from a malaria endemic region.

## Material and methods

### Study design and study population

The samples (n=67) analyzed here were part of a case-control study of severe malaria conducted in a region of intense malaria transmission in Northern Uganda. The demographic and clinical details of the study subjects recruited into the study at Apac Hospital have been reported (18). Briefly, prior to the start of the study, ethical approval was obtained from the Uganda National Council for Science and Technology and written parental consent was obtained prior to the enrolment of the children. The study population were children aged 6-59 months old who were attending or were admitted at Apac Hospital with a primary diagnosis of malaria. The controls, matched for age, sex, and geographic location or parish residence, were recruited from the Outpatient Department when they presented with other illnesses and were well enough to go home. Children were excluded when they had life threatening illnesses and clinical signs of pneumonia, bacterial or parasite-related gastroenteritis, signs of HIV-1/AIDS-related opportunistic infections, or their parents/guardians did not provide informed consent. Two-ml venous blood samples were collected from both cases and controls; serum samples were stored at −20°C or −80°C until analyzed. Giemsa-stained thin blood smears were microscopically examined to determine parasite density and pigmented monocytes. Hemoglobin levels and packed cell volumes were measured colorimetrically and using a hematocrit centrifuge, respectively.

For this study, children with severe and complicated malaria (SM) were those hospitalized with fever and *P. falciparum* asexual parasitaemia of above ≥5,000 parasites/*μ*L and at least one indicator of severe disease: severe malarial anaemia (< 5 g/dL), respiratory distress (deep breathing), prostration, and/or convulsions. Children were excluded if they had septicemia, typhoid fever, pylonephritis, lobar pneumonia, and viral hepatitis. Malaria control subjects with uncomplicated malaria (UM) were children with either a positive parasitemia of ≥5,000 parasites/μL and body temperatures ≥37.5 °C but did not manifest any sign of complicated malaria, or children with a positive parasitemia of ≤ 5,000 parasites/μL but who had neither fever nor any signs of complicated disease (AS; asymptomatic malaria).

### Enzyme-linked immunosorbent assay

HMGB1 levels in sera of individual children from each clinical sub-category were determined by quantitative enzyme-linked immunosorbent assay kits according to the manufacturer’s recommendations (SHINO-TEST Corporations, Kanagawa, Japan). Briefly, each reaction contained specific mouse anti-human HMGB1 capture monoclonal antibody, a reporter anti-mouse polyclonal antibody conjugated to horseradish peroxidase. Individual serum samples were 100-fold diluted, assayed in duplicate, and levels of specific factors determined spectrophotometrically by an ELISA reader at 450 nm using a standard curve prepared from human HMGB1 protein, which was also used as positive control.

### Statistical methods

Data analysis was performed using The R software (R version 4.0.1 - “See Things Now”; R Foundation for Statistical Computing). The differences in serum levels of chemokines in children with severe, complicated and uncomplicated malaria were assessed using the Kruskal-Wallis test (*p*<0.001); with Dunn’s test as post hoc test. Statistical difference between genders and pigmentation status was assessed using the Mann-Whitney U test. The correlation between HMGB1 levels and markers of malaria morbidity or immunity was assessed by the Spearman’s correlation coefficient test. To determine independent correlates of serum HMGB1 levels, multiple stepwise linear regression analysis was performed. Mean levels of serum HMGB1 stratified by the independent correlates were then compared using analysis of covariance (ANCOVA), adjusted for age and sex. Statistical significance was defined as *p* < 0.05.

## Results

### Characteristics of study participants

The objective of this study was to measure serum concentrations of HMGB1 and enumerate pigmented monocytes in female and male infants with different clinical malaria outcomes to determine if these parameters differed by sex. These infants constituted a subset of study participants from a previously published study of severe malaria (18). Demographic data for this subset of participants are presented in Table 1. Infants were categorized as having SM, UM, AS or being healthy controls (HC) as defined in Materials and Methods. The median (and interquartile range; IQR) age was similar in infants with the different malaria outcomes and in female and male infants.

**Table 1:**
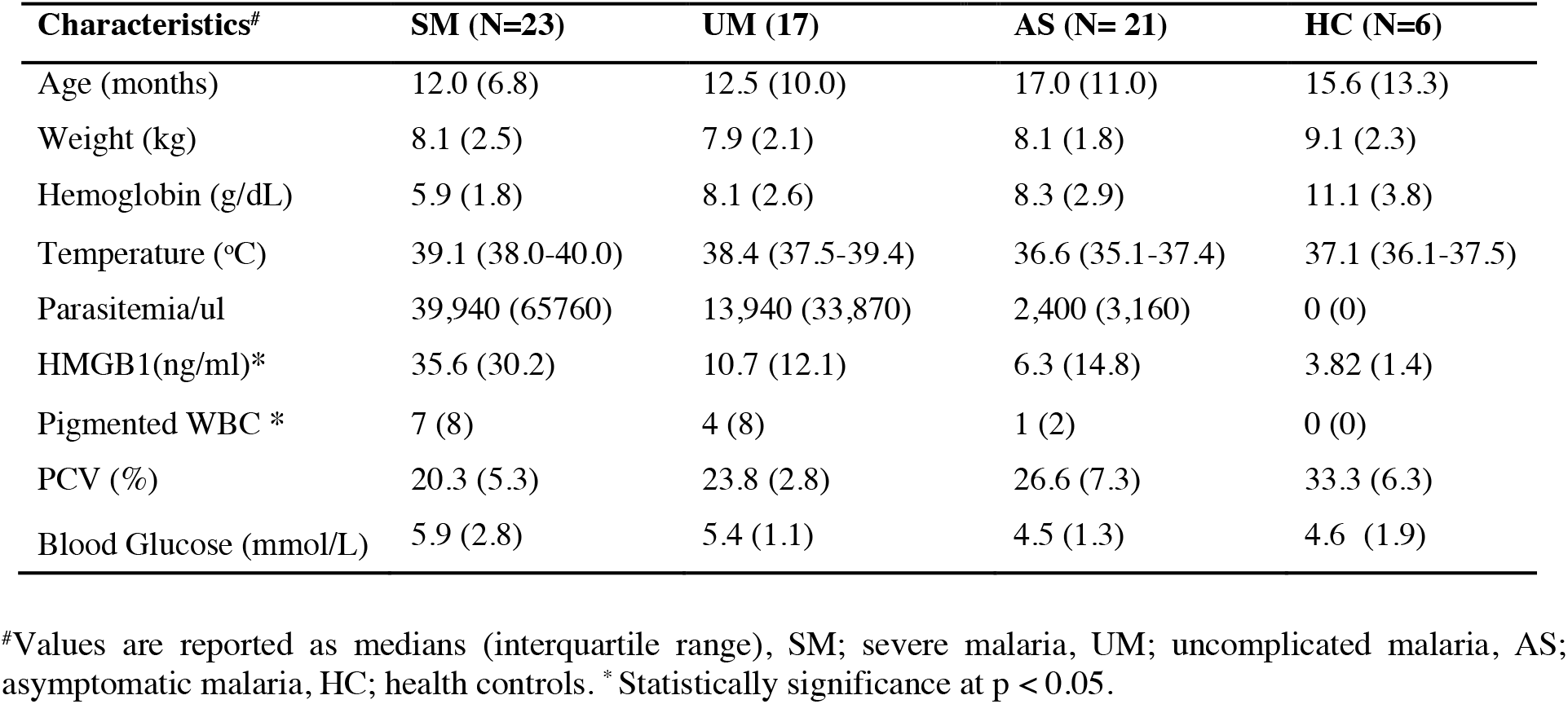
Characteristics of subjects assessed in this study

| Characteristics<sup>#</sup> | SM (N=23) | UM (17) | AS (N= 21) | HC (N=6) |
| --- | --- | --- | --- | --- |
| Age (months) | 12.0 (6.8) | 12.5 (10.0) | 17.0 (11.0) | 15.6 (13.3) |
| Weight (kg) | 8.1 (2.5) | 7.9 (2.1) | 8.1 (1.8) | 9.1 (2.3) |
| Hemoglobin (g/dL) | 5.9 (1.8) | 8.1 (2.6) | 8.3 (2.9) | 11.1 (3.8) |
| Temperature (°C) | 39.1 (38.0-40.0) | 38.4 (37.5-39.4) | 36.6 (35.1-37.4) | 37.1 (36.1-37.5) |
| Parasitemia/ul | 39,940 (65760) | 13,940 (33,870) | 2,400 (3,160) | 0 (0) |
| HMGB1(ng/ml)* | 35.6 (30.2) | 10.7 (12.1) | 6.3 (14.8) | 3.82 (1.4) |
| Pigmented WBC * | 7 (8) | 4 (8) | 1 (2) | 0 (0) |
| PCV (%) | 20.3 (5.3) | 23.8 (2.8) | 26.6 (7.3) | 33.3 (6.3) |
| Blood Glucose (mmol/L) | 5.9 (2.8) | 5.4 (1.1) | 4.5 (1.3) | 4.6 (1.9) |

<sup>#</sup>Values are reported as medians (interquartile range), SM; severe malaria, UM; uncomplicated malaria, AS; asymptomatic malaria, HC; health controls. \*Statistically significance at $p < 0.05$ .

### Sex differences in serum HMGB1 concentrations

Serum concentrations of HMGB1 in infants with different malaria outcomes were determined by ELISA (Fig. 1A). The median (and IQR) HMGB1 concentration was significantly higher in SM (35.6 (30.2) ng/ml) than in UM (10.7 (12.1) ng/ml; *p*=0.031) and AS patients (6.3 (14.8) ng/ml; *p*<0.01). Age-matched HC had significantly (*p*=0.001) lower HMGB1 concentrations compared to malaria-infected infants in each of the three clinical categories. Compared with HC and AS infants, infants with clinical malaria (UM and SM) showed significantly higher HMGB1 concentrationss (*p*<0.001). However, when serum HMGB1 concentrations were normalized by dividing with the number of pigmented monocytes per infant, AS infants showed an unexpected trend towards higher HMGB1 levels by comparison with infants with clinical malaria (UM, SM; *p*=0.06; Fig S1A).

**Figure 1:**
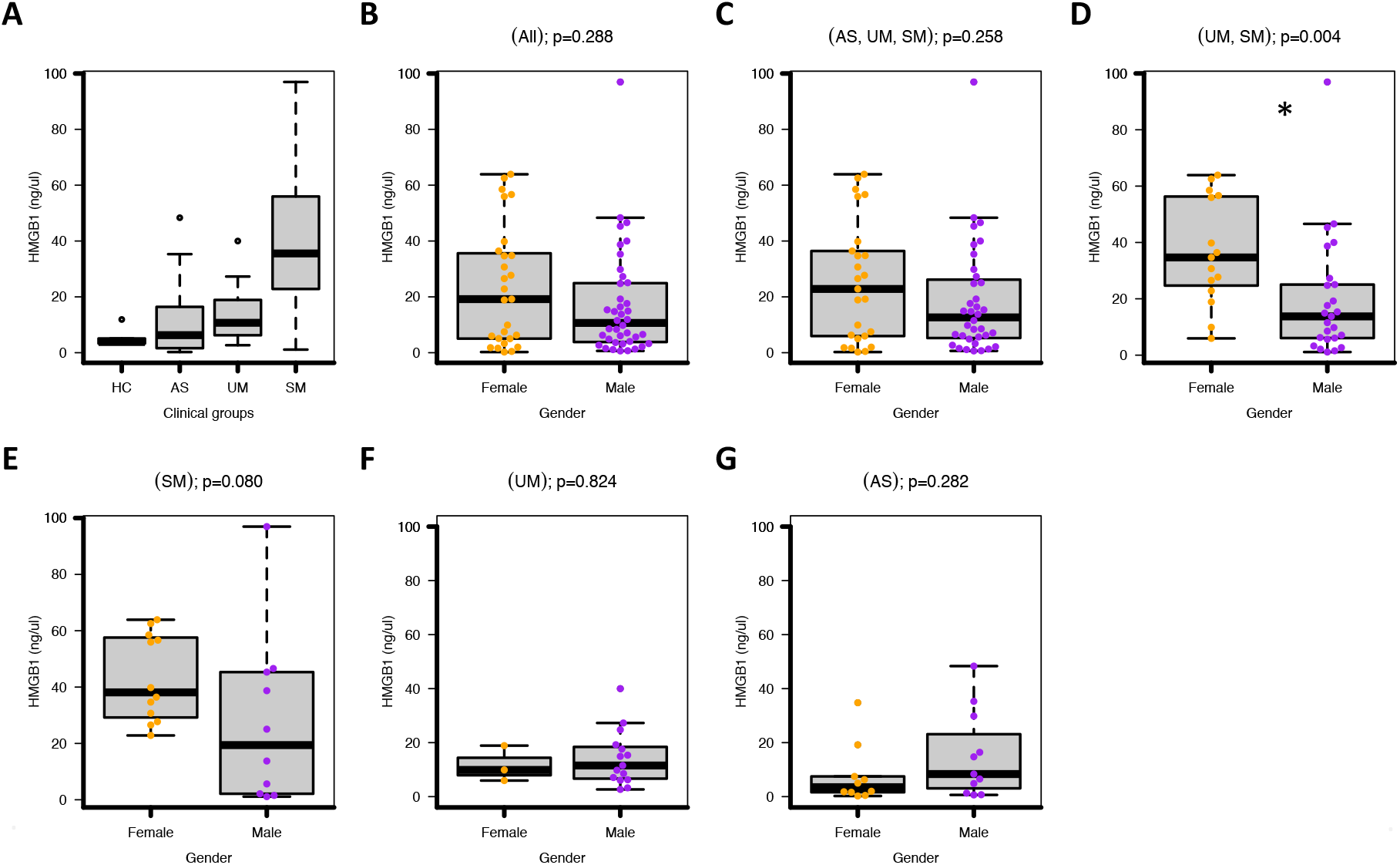
Levels of HMGB1 by clinical groups and gender. **A**: HMGB1 levels as determined by sandwich ELISA for 23 patients with SM, 17 with UM, and 21 with AS, alongside 6 healthy controls (HC) are shown. The horizontal lines in the box represent the medians of the distributions. Statistical difference was assessed using the Kruskal-Wallis test (*P* < 0.001); with Dunn’s test as post hoc test, *P* < 0.01. **B-G**: HMGB1concentrations in individual female and male are shown, with box-plots indicating the median and ranges for each age cohort. Statistical difference between genders was assessed using the Mann-Whitney U test. (B) All children, (C) AS, UM and SM, (D) UM and SM (E) only SM, (F) only UM and (F) only AS. * Statistical difference was observed for UM and SM clinical groups at p=0.004.

We then compared serum concentrations of HMGB1 in male and female infants. We did not observe any difference by sex when all infants were considered irrespective of malaria outcomes (Fig 1B). Sex differences only emerged when different malaria outcomes were considered separately. Specifically, median concentrations of HMGB1 were comparable in malaria-infected (SM, UM, and AS combined) male and female infants (Fig 1C). However, in infants with clinical malaria (UM and SM combined) females had significantly higher concentrations of HMGB1 compared with males (*p*=0.004; Fig ID). When the HMGB1 concentrations were normalized, female infants showed a trend towards higher HMGB1 concentrations per pigmented monocyte albeit not significant (*p*=0.097; Fig S1B). The lack of significance after normalization may suggest that other innate immune cells or cellular mechanisms could be contributing to HMGB1 levels in clinical malaria. When SM was considered alone, there was a trend for higher HMGB1 concentrations in females but this difference was marginally nonsignificant (*p*=0.08; Fig. 1E) for non-normalized concentrations and nonsignificant for normalized concentrations (*p*=0.133; Fig. S1C). When UM was considered alone, female and male infants had comparable nonnormalized serum HMGB1 concentrations (*p*=0.82; Fig 1F). In infants with AS, males showed a trend for higher nonnormalized and normalization HMGB1 concentrations compared to females but this was not significant (Fig 1G and Fig S1D).

### Sex differences in counts of pigmented monocytes

Giemsa-stained thin blood smears prepared from malaria-infected patients were examined microscopically for the enumeration of monocytes with hemozoin, the biocrystal malaria pigment synthesized by *Plasmodium* spp. to avoid the toxicity of free heme after digestion of hemoglobin (19). Numbers of pigmented monocytes were significantly different between various malaria outcomes (Fig 2A). When infants were stratified by sex, the numbers of pigmented monocytes were comparable between females and males for all infants combined irrespective of malaria infection status, for all malaria-infected infants (AS, UM, SM), for infants with clinical malaria (UM, SM combined), and for infants with SM alone (Fig 2B-F). However, in infants with AS, males had significantly higher numbers of pigmented monocytes than females (*p*=0.021; Fig 2G).

**Figure 2:**
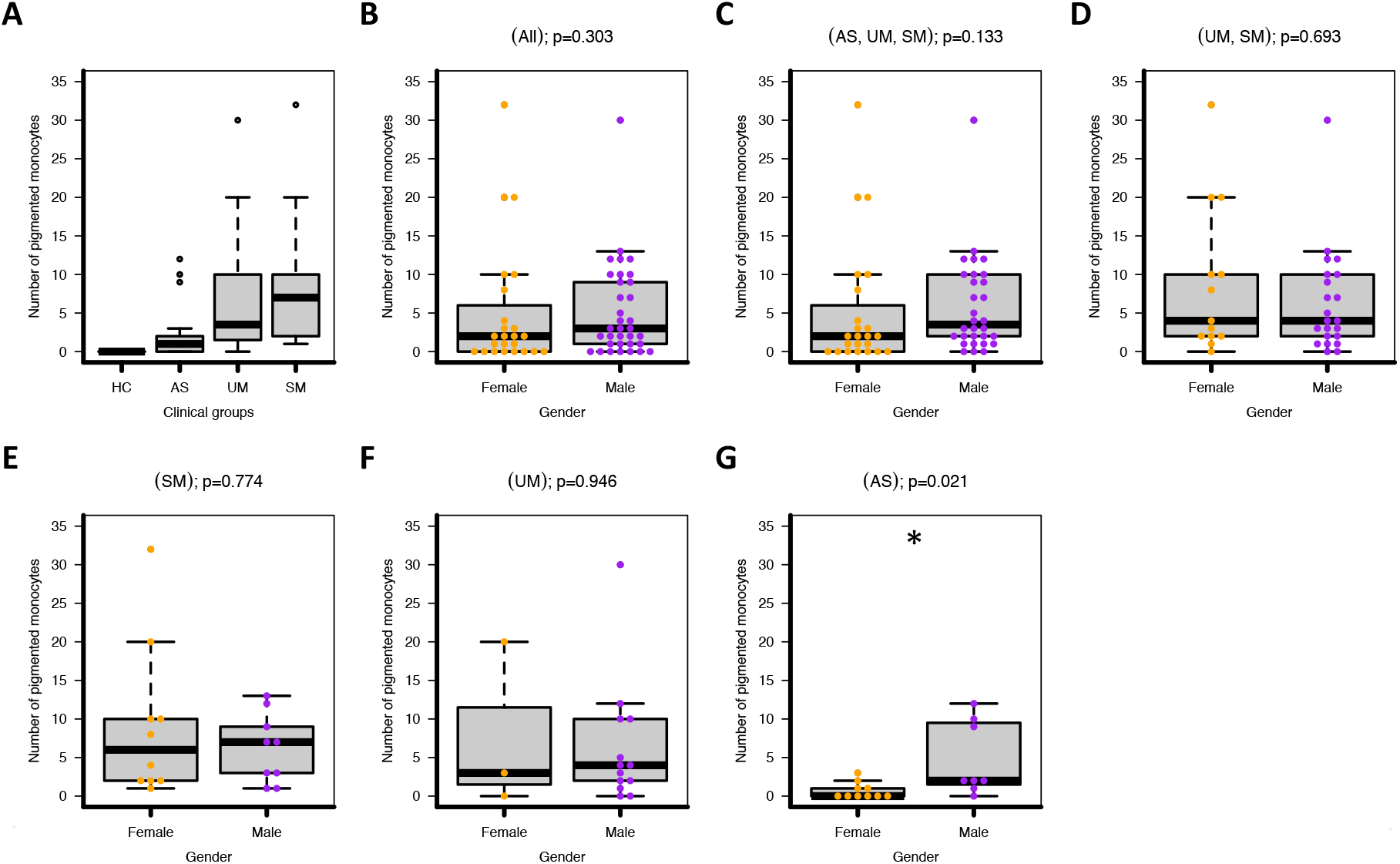
Number of pigmented monocytes by clinical groups and gender. **A:** Number of pigmented monocytes by clinical groups is shown. The horizontal lines in the box-plots represent the medians of the distributions. Statistical difference was assessed using the Kruskal-Wallis test (P < 0.001); with Dunn’s test as post hoc test, P < 0.01. **B-G.** Number of pigmented monocytes in different genders by clinical outcome is shown with box-plots indicating the median and ranges for each age cohort. Statistical difference between genders was assessed using the Mann-Whitney U test. (B) All children, (C) AS, UM and SM, (D) UM and SM (E) only SM, (F) only UM and (F) only AS. * Statistical difference was observed in the AS malaria clinical at p=0.021.

### Parasite density in male and female infants

Since the activation of monocytes/macrophages to take up hemozoin and produce HMGB1 probably depend on parasite density in individual patients, we compared parasite density between both sexes. Female infants showed a trend towards higher parasite densities compared to males (Fig S2A-C) with a significant difference being observed among the clinical groups (UM, SM; Fig S2B *p*=0.019).

### Correlations between HMGB1, pigmented monocyte and parasite density

We investigated whether there were any significant correlations between parasite density and serum levels of HMGB1 and numbers of pigmented monocyte in all female and male children. First, we observed a significant correlation between numbers of pigmented monocytes and concentrations of HMGB1 but this was significant only in female (r=0.40, *p*=0.043) but not in male infants (r= 0.20, *p*=0.214) (Fig. 3A). Second, there was a significant correlation between parasite density and concentrations of HMGB1 in female (r=0.51, *p*=0.039) but not male infants (r= −0.02, *p*=0.667) (Fig 3B). Finally, there was a significant correlation between parasite density and numbers of pigmented monocytes in female (r=0.44, *p*=0.011) but not male infants (r=0.14, *p*=0.377) (Fig 3C).

**Figure 3:**
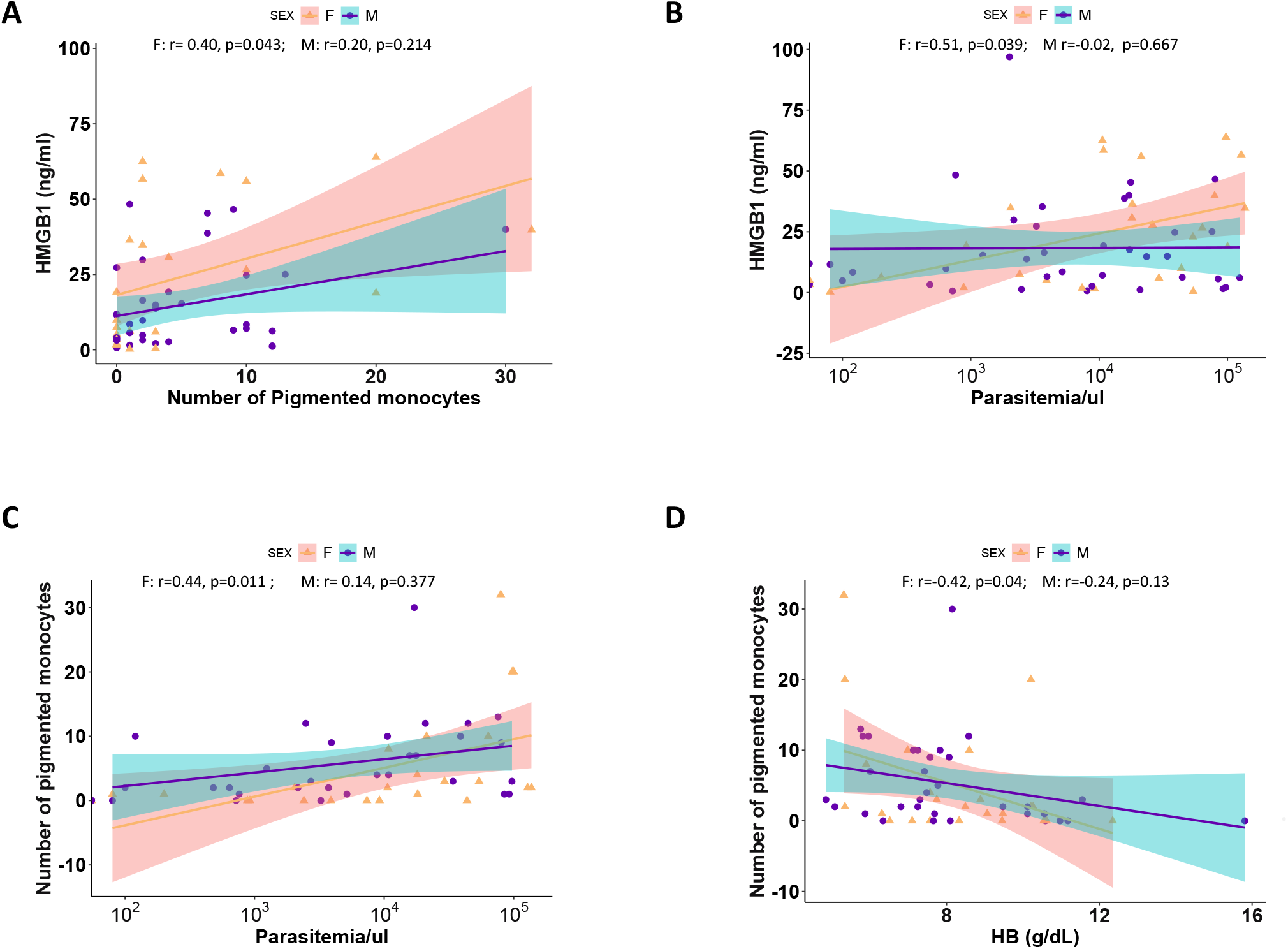
Correlation between Serum HMGB1 levels and markers of malaria morbidity. Pearson correlation coefficients (R) and p values for each sex are shown. Lines represent linear regression lines and shading represents 95% confidence intervals. **A.** HMGB1 vs number of pigmented monocytes. **B.** HMGB1 vs parasitemia. **C**. Number of pigmented monocytes vs parasitemia. **D**. Number of pigmented monocytes vs hemoglobin levels.

### Correlations between HMGB1, pigmented monocyte and hemoglobin

We then assessed whether there were any correlations between hemoglobin (HB) concentration as a proxy marker of immunity and serum concentrations of HMGB1 and numbers of pigmented monocyte in female and male children. HB concentrations, as expected, demonstrated a negative correlation with pigmented monocytes in female (r=0.42, *p*=0.041) but not male infants (r=0.24, *p*=0.133) (Fig 3D). However, there was no correlation between serum HMGB1concentartions and HB (Supplementary Fig S3). Correlations between HB, age, packed cell volumes and other infant parameters compared by gender are summarized in Supplementary Fig S3.

## Discussion

Severe malaria in African infants is characterized by high serum concentrations of HMGB1 (17) and increased numbers of pigmented monocytes (14, 18). HMGB1, passively released by dead or dying cells as a result of tissue injury following sterile inflammation or infection with pathogens, serves as a danger signal (12). As a damage associated molecular pattern (DAMP) molecule, extracellular HMGB1 excerbates inflammatory responses by activating monocytes/macrophages to produce proinflammatory cytokines (20) that characterize severe disease in sepsis (15), COVID-19 (13), and severe and complicated malaria (14). HMGB1 is also actively secreted by activated monocytes/macrophages (16). It is not known which mechanism accounts for serum concentrations of HMGB1 in infants with malaria. Pigmented monocytes on the other hand are activated monocytes which have phagoctosed hemozoin or malaria pigment produced when malaria parasites convert haem released by haemoglobin digestion (21). The relationship between pigmented monocytes and other biomarkers of morbidity like HMGB1 in infants with malaria has hitherto remained unknown.

The present study shows preliminary results of what appears to be important sex differences in serum concentrations of HMGB1 and numbers of pigmented monocytes in infants with asymptomatic and clinical malaria. First, male and female infants with clinical malaria had comparable numbers of pigmented monocytes but female infants had significantly higher serum concentrations of HMGB1. These data suggested that female infants produced more HMGB1 than male infants. This was confirmed when serum concentrations of HMGB1 were normalized to numbers of pigmented monocytes in each infant. Second, male and female infants with asymptomatic malaria had comparable serum concentrations of HMGB1 but male infants had significantly higher numbers of pigmented monocytes. High numbers of pigmented monocytes in males did not therefore translate into high HMGB1 serum levels. The fact that asymptomatic malaria is characterized by lack of clinical symptoms, antiinflammatory cytokines, and higher numbers of pigmented monocytes in males could suggest that the monocyte/macrophage subset in asymptomatic malaria might be distinct from that in clinical malaria and does not produce significant quantities of HMGB1. We speculate that these monocytes are of the M2 phenotype which produce anti-inflammatory cytokines (22) that maintain anti-disease immunity by dampening the proinflammatory cytokine response and HMGB1 production. This would explain the comparable serum HMGB1 concentrations in males and female infants despite significantly elevated numbers of pigmented monocytes in males.

Further we carried out correlation analyses in order to understand the relationships between parasite density as a marker of morbidity and both HMGB1 and pigmented monocytes.

Although parasite density was comparable in male and female infants the correlation between HMGB1, pigmented monocytes and parasite density was significant only in female infants. This suggests that *P. falciparum* parasites or parasite products trigger signalling pathways in innate immune cells from females but not males. It is known that the necrotic or necroptotic pathway of cell death which results in the release of HGMB1 by dying cells occurs only in females (23). The relevance of this mechanism to our findings in Ugandan infants with malaria remains to be determined. We also investigated the correlation between hemoglobin as a proxy marker of immunity in infants with malaria and pigmented monocytes. Numbers of pigmented monocytes negatively correlated with haemoglobin levels and this correlation was significant only in female but not male infants. This underscores the fact that female infants are at a greater risk of monocyte/macrophage-mediated malaria morbidity exacerbated by the higher production of HMGB1 by monocytes/macrophages in female infants with clinical malaria.

Sex differences in innate and adaptive immunity can be attributed to the effect of sex hormones or genetic differences between females and males (4). Since our study participants were infants < 5 years old, well below the age of puberty, the effect of sex hormones is minimal. Sex differences in immune responses between pre-pubertal female and male children have been reported in other studies which strongly suggest that genetic differences are at play (4, 9, 24, 25). We propose therefore that genetic differences may account for the observed sexual dimorphism in serum HMGB1 concentrations and numbers of pigmented monocytes in this population.

This study has several limitations. First, the case-control approach used in the original study does not allow hypotheses to be generated about possible cause-and effect relationships. Second, the small sample sizes within different malaria outcomes probably prevented us from observing significant differences or overestimated others. Third, flow cytometric characterization of monocytes/macrophages by CD14 and CD16 surface markers to establish functional differences in monocyte/macrophage populations in clinical and asymptomatic malaria was not feasible at the time of the study. Fourth, the role of cell types other than monocytes/macrophages such as neutrophils, which constituted 1 % of pigmented leucocytes in the study population (18), was not investigated. These limitations notwithstanding, the sex differences in innate immunity involving pigmented monocytes and HMGB1 in infants with asymptomatic and clinical malaria are markedly different and further longitudinal studies would address these limitations.

In conclusion, our study has presented preliminary evidence of sex differences in innate immune responses involving HMGB1 and pigmented monocytes during clinical and asymptomatic malaria in Ugandan infants <5 years old residents of a malaria-endemic region. The study provides important preliminary insights into possible mechanisms underlying the higher risk of morbidity in female infants with clinical malaria. If confirmed across various geographic regions of different malaria endemicities, the data may inform the development of sex-based therapeutic approaches which minimize malaria-related morbidity and mortality in females while being equally effective in both sexes. It is noteworthy that specific anti-HMGB1 therapeutics have been proposed for the treatment of severe COVID-19 (26) and sepsis (27) which are similar to severe malaria in monocyte/macrophage-mediated morbidity. Put together, these findings suggest that female infants with clinical malaria might be at a greater risk of morbidity characterized by higher serum HMGB1 levels and call for further studies.

## Acknowledgments

We appreciate the study volunteers at Apac Hospital, Northern Uganda; and thank the research team from Med Biotech Laboratories and at the Apac Hospital for their technical assistance in obtaining the study samples. We thank Prof. Nirianne M. Q. Palacpac of Osaka University for critical reading of the manuscript.

## Supplementary Figures

**Figure S1:**
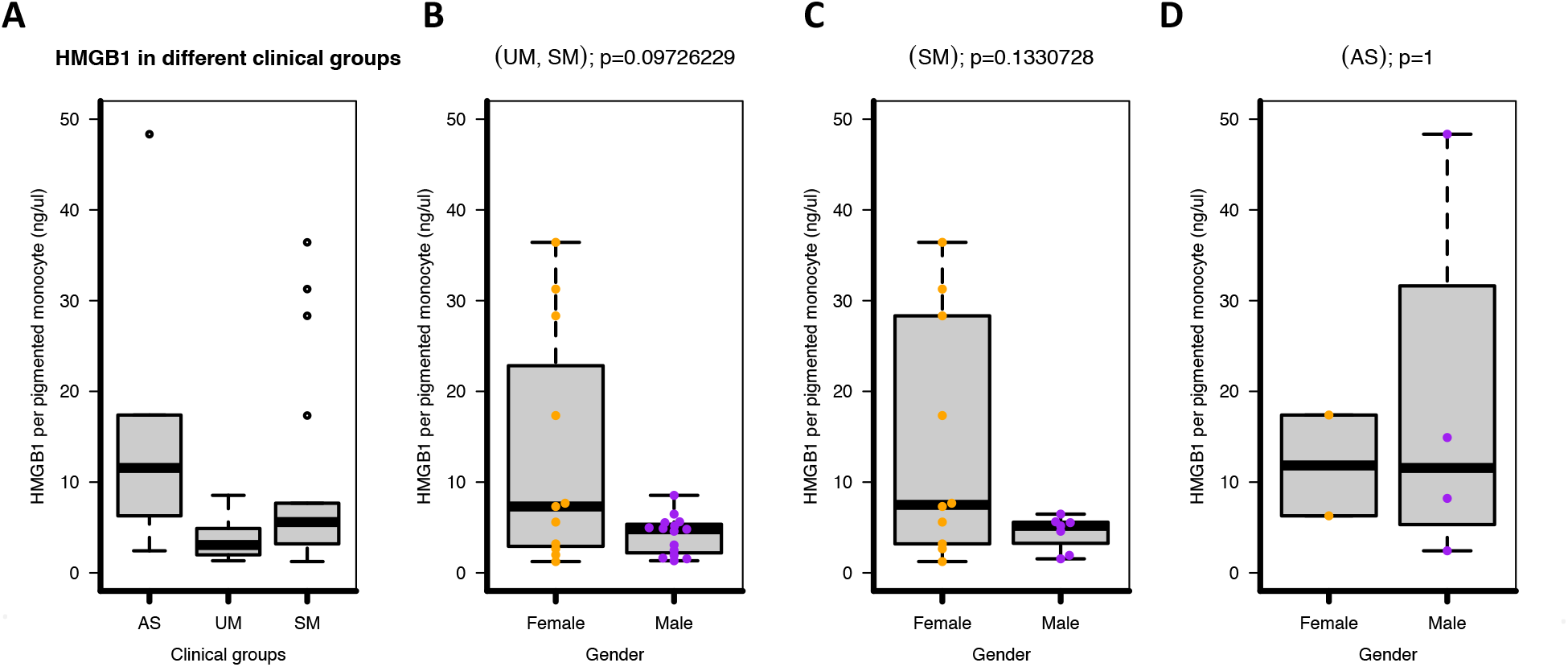
Normalized HMGB1 per pigmented monocyte. Serum HMGB1 concentrations were normalized by dividing with the number of pigmented monocytes per infant **A**: HMGB1 per pigmented monocyte by clinical groups is shown. Statistical difference was assessed using the Kruskal-Wallis test (*P* < 0.01); with Dunn’s test as post hoc test, *P* < 0.01. **B-C**: HMGB1 per pigmented monocyte in different genders by clinical outcome is shown. Statistical difference was assessed using the Mann-Whitney U test. (B) UM and SM, (C) only SM, and (D) only AS.

**Figure S2:**
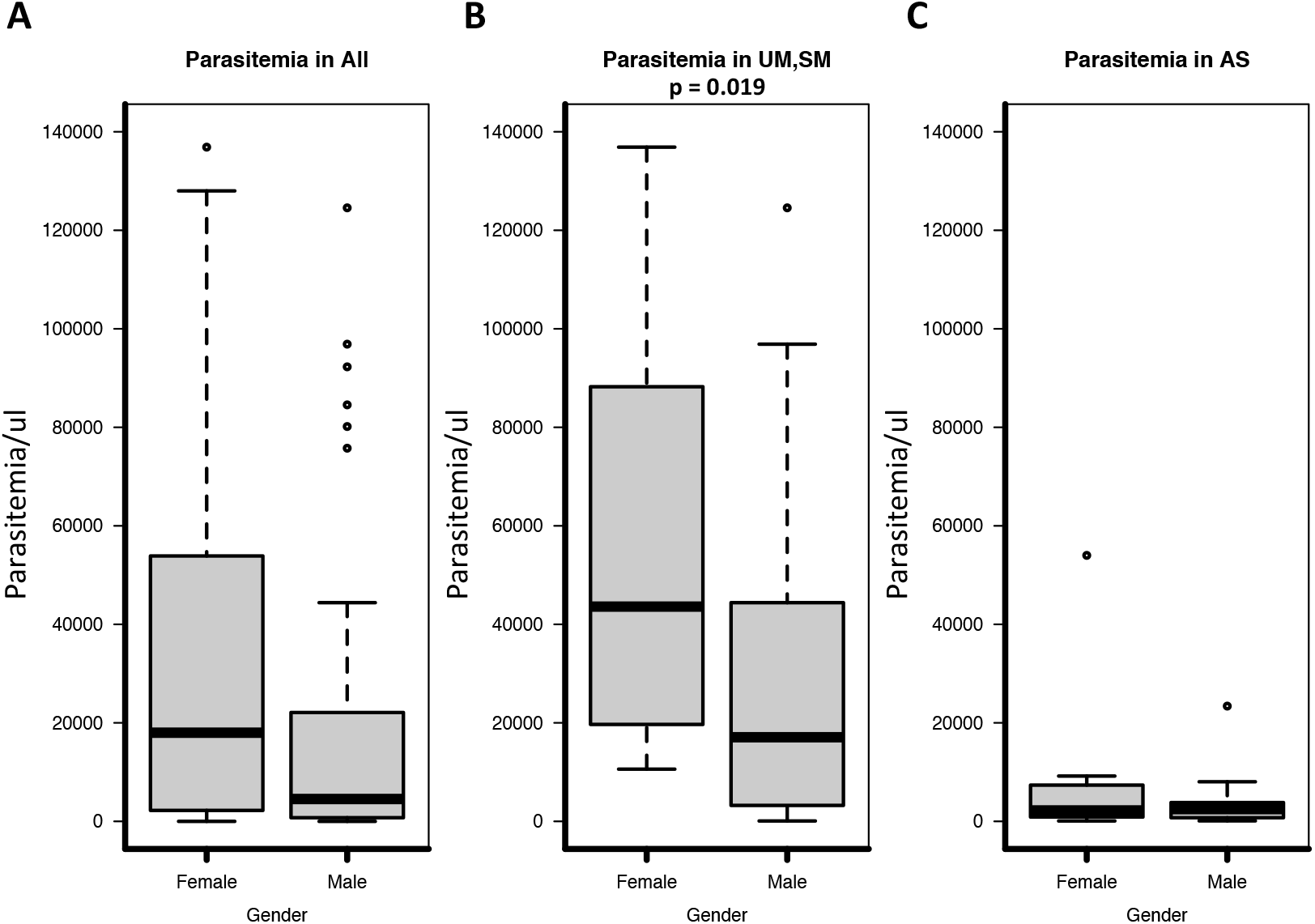
Parasite density by gender. The horizontal lines in the box-plots represent the medians of the distributions. Statistical difference between genders was assessed using the Mann-Whitney U test. **A**. All participants (p > 0.05). **B**. In the UM and SM group (p = 0.019) **C.** In the AS group only (p > 0.05).

**Figure S3:**
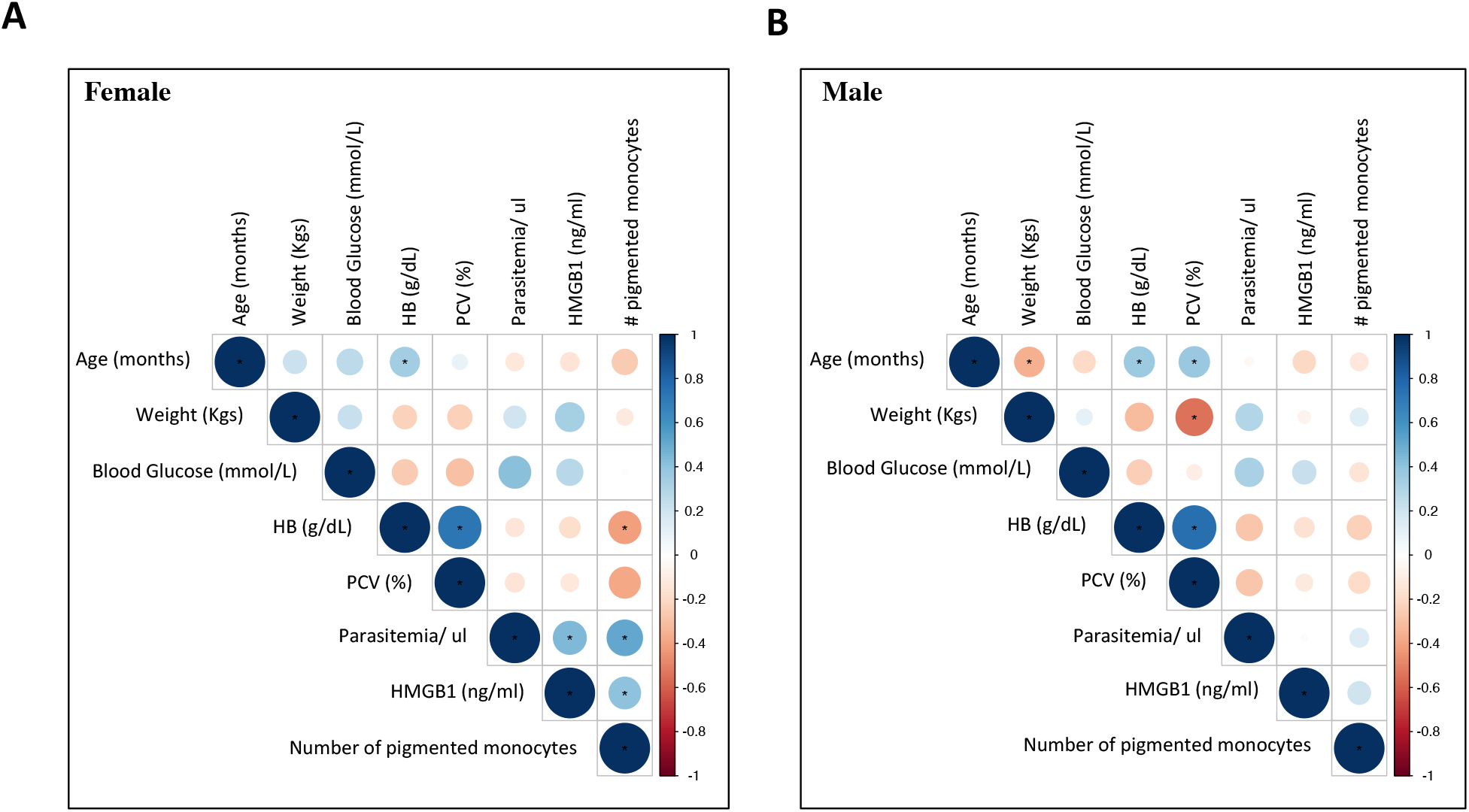
Heatmap showing Pearson correlation of HMGB1, pigmented monocyte and markers of malaria morbidity for each gender. The size and colour of the circles indicate the correlation coefficient (R). * indicates statistically significant correlations (p < 0.05).

## References

1. WHO. (2019) World Malaria Report 2019. p. 232, World Health Organization, Geneva, Switzerland, Geneva, Switzerland

2. S, K., P, K. T., B, N. M., N, M., Cbk, M., and Karuna, T. (2014) Severe Falciparum Malaria-Difference in Mortality among Male and Nonpregnant Females. Journal of clinical and diagnostic research : JCDR 8, MC01–04

3. Fischinger, S., Boudreau, C. M., Butler, A. L., Streeck, H., and Alter, G. (2019) Sex differences in vaccine-induced humoral immunity. Semin. Immunopathol. 41, 239–249

4. Klein, S. L., and Flanagan, K. L. (2016) Sex differences in immune responses. Nat Rev Immunol 16, 626–638

5. Okech, B. A., Corran, P. H., Todd, J., Agwang, C., Riley, E. M., and Egwang, T. G. (2020) Differences in Anti Malarial Antibody Concentrations and Specificities between Male and Female Ugandan Children. Archives of Immunology 2, 11–17

6. Migasena, S., Heppner, D. G., Kyle, D. E., Chongsuphajaisiddhi, T., Gordon, D. M., Suntharasamai, P., Permpanich, B., Brockman, A., Pitiuttutham, P., Wongsrichanalai, C., Srisuriya, P., Phonrat, B., Pavanand, K., Viravan, C., and Ballou, W. R. (1997) SPf66 malaria vaccine is safe and immunogenic in malaria naive adults in Thailand. Acta Trop 67, 215–227

7. Nosten, F., Luxemburger, C., Kyle, D. E., Gordon, D. M., Ballou, W. R., Sadoff, J. C., Brockman, A., Permpanich, B., Chongsuphajaisiddhi, T., and Heppner, D. G. (1997) Phase I trial of the SPf66 malaria vaccine in a malaria-experienced population in Southeast Asia. Am J Trop Med Hyg 56, 526–532

8. Klein, S. L., Shann, F., Moss, W. J., Benn, C. S., and Aaby, P. (2016) RTS,S Malaria Vaccine and Increased Mortality in Girls. mBio 7, e00514–00516

9. Prahl, M., Jagannathan, P., McIntyre, T. I., Auma, A., Wamala, S., Nalubega, M., Musinguzi, K., Naluwu, K., Sikyoma, E., Budker, R., Odorizzi, P., Kakuru, A., Havlir, D. V., Kamya, M. R., Dorsey, G., and Feeney, M. E. (2017) Sex Disparity in Cord Blood FoxP3(+) CD4 T Regulatory Cells in Infants Exposed to Malaria In Utero. Open Forum Infect Dis 4, ofx022

10. Briggs, J., Teyssier, N., Nankabirwa, J. I., Rek, J., Jagannathan, P., Arinaitwe, E., Bousema, T., Drakeley, C., Murray, M., Crawford, E., Hathaway, N., Staedke, S. G., Smith, D., Rosenthal, P. J., Kamya, M., Dorsey, G., Rodriguez-Barraquer, I., and Greenhouse, B. (2020) Sex-based differences in clearance of chronic Plasmodium falciparum infection. Elife 9

11. Ortega-Pajares, A., and Rogerson, S. J. (2018) The Rough Guide to Monocytes in Malaria Infection. Front. Immunol. 9, 2888

12. Venereau, E., Ceriotti, C., and Bianchi, M. E. (2015) DAMPs from Cell Death to New Life. Front. Immunol. 6, 422

13. Costela-Ruiz, V. J., Illescas-Montes, R., Puerta-Puerta, J. M., Ruiz, C., and Melguizo-Rodriguez, L. (2020) SARS-CoV-2 infection: The role of cytokines in COVID-19 disease. Cytokine Growth Factor Rev. 54, 62–75

14. Lyke, K. E., Burges, R., Cissoko, Y., Sangare, L., Dao, M., Diarra, I., Kone, A., Harley, R., Plowe, C. V., Doumbo, O. K., and Sztein, M. B. (2004) Serum levels of the proinflammatory cytokines interleukin-1 beta (IL-1beta), IL-6, IL-8, IL-10, tumor necrosis factor alpha, and IL-12(p70) in Malian children with severe Plasmodium falciparum malaria and matched uncomplicated malaria or healthy controls. Infect Immun 72, 5630–5637

15. Matsumoto, H., Ogura, H., Shimizu, K., Ikeda, M., Hirose, T., Matsuura, H., Kang, S., Takahashi, K., Tanaka, T., and Shimazu, T. (2018) The clinical importance of a cytokine network in the acute phase of sepsis. Sci Rep 8, 13995

16. Gardella, S., Andrei, C., Ferrera, D., Lotti, L. V., Torrisi, M. R., Bianchi, M. E., and Rubartelli, A. (2002) The nuclear protein HMGB1 is secreted by monocytes via a non-classical, vesicle-mediated secretory pathway. EMBO Rep 3, 995–1001

17. Higgins, S. J., Xing, K., Kim, H., Kain, D. C., Wang, F., Dhabangi, A., Musoke, C., Cserti-Gazdewich, C. M., Tracey, K. J., Kain, K. C., and Liles, W. C. (2013) Systemic release of high mobility group box 1 (HMGB1) protein is associated with severe and fatal Plasmodium falciparum malaria. Malaria journal 12, 105

18. Mujuzi, G., Magambo, B., Okech, B., and Egwang, T. G. (2006) Pigmented monocytes are negative correlates of protection against severe and complicated malaria in Ugandan children. Am J Trop Med Hyg 74, 724–729

19. Pagola, S., Stephens, P. W., Bohle, D. S., Kosar, A. D., and Madsen, S. K. (2000) The structure of malaria pigment beta-haematin. Nature 404, 307–310

20. Bianchi, M. E., Crippa, M. P., Manfredi, A. A., Mezzapelle, R., Rovere Querini, P., and Venereau, E. (2017) High-mobility group box 1 protein orchestrates responses to tissue damage via inflammation, innate and adaptive immunity, and tissue repair. Immunol. Rev. 280, 74–82

21. Coronado, L. M., Nadovich, C. T., and Spadafora, C. (2014) Malarial hemozoin: from target to tool. Biochim. Biophys. Acta 1840, 2032–2041

22. Viola, A., Munari, F., Sanchez-Rodriguez, R., Scolaro, T., and Castegna, A. (2019) The Metabolic Signature of Macrophage Responses. Front. Immunol. 10, 1462

23. Zemskova, M., Kurdyukov, S., James, J., McClain, N., Rafikov, R., and Rafikova, O. (2020) Sex-specific stress response and HMGB1 release in pulmonary endothelial cells. PLoS One 15, e0231267

24. Casimir, G. J., Lefevre, N., Corazza, F., and Duchateau, J. (2013) Sex and inflammation in respiratory diseases: a clinical viewpoint. Biol. Sex Differ. 4, 16

25. Voysey, M., Pollard, A. J., Perera, R., and Fanshawe, T. R. (2016) Assessing sex-differences and the effect of timing of vaccination on immunogenicity, reactogenicity and efficacy of vaccines in young children: study protocol for an individual participant data meta-analysis of randomised controlled trials. BMJ Open 6, e011680

26. Andersson, U., Ottestad, W., and Tracey, K. J. (2020) Extracellular HMGB1: a therapeutic target in severe pulmonary inflammation including COVID-19? Mol. Med. 26, 42

27. Stevens, N. E., Chapman, M. J., Fraser, C. K., Kuchel, T. R., Hayball, J. D., and Diener, K. R. (2017) Therapeutic targeting of HMGB1 during experimental sepsis modulates the inflammatory cytokine profile to one associated with improved clinical outcomes. Sci Rep 7, 5850

